# Multi-omics analysis identifies essential regulators of mitochondrial stress response in two wild-type *C. elegans* strains

**DOI:** 10.1101/2021.07.20.453059

**Authors:** Arwen W. Gao, Gaby El Alam, Amélia Lalou, Terytty Yang Li, Marte Molenaars, Yunyun Zhu, Katherine A. Overmyer, Evgenia Shishkova, Kevin Hof, Maroun Bou Sleiman, Riekelt H. Houtkooper, Joshua J. Coon, Johan Auwerx

## Abstract

The mitochondrial unfolded protein response (UPRmt) is a promising pharmacological target for aging and age-related diseases. However, the integrative analysis of the impact of UPRmt activation on different layers of signaling in animals with a different genetic background is lacking. In this study, we applied systems approaches to investigate the effect of UPRmt induced by administering doxycycline (Dox) on transcriptome, proteome, lipidome, and metabolome in two genetically divergent *C. elegans* strains. We found that Dox prolongs lifespan of both worm strains through pathways in both shared and strain-specific manners. From the integrated omics datasets, we observed a strong impact of Dox on mitochondrial functions, detected upregulated defense response and lipid metabolism, identified decreased triglycerides and lowered metabolome profiles in both strains. This conserved phenomic footprint has great translational value as it indicates that the beneficial effects of Dox-induced UPRmt on health and lifespan are consistent across different genetic backgrounds.

## Introduction

Mitochondria are essential organelles for numerous processes, such as energy harvesting, intermediate metabolism, autophagy, and immune response (Nunnari and Suomalainen 2012; Quiros et al. 2016; West and Shadel 2017). Changes in mitochondrial number, morphology, and functions not only impact cellular metabolism but also influence whole body metabolism, health, and lifespan (Nunnari and Suomalainen 2012; Vafai and Mootha 2012; Andreux et al. 2013). There has been increasing evidence that mitochondrial dysfunction accumulates upon aging and correlates with the development of many age-associated diseases (Sun et al. 2016). Age-associated mitochondrial impairments include decreased efficiency of oxidative phosphorylation, increases in oxidative damage, aggregation of mitochondrial proteins, alteration of mitochondrial quality control (e.g. mitophagy), accumulation of mtDNA mutations, as well as dysregulation of many aspects of mitochondrial metabolism (Sun et al. 2016; Jang et al. 2018; D’Amico et al. 2019).

Given the central role of mitochondria in health- and lifespan, mitochondria evolved an elaborate quality control system directing pleiotropic mitochondrial stress response (MSR) pathways to ensure optimal mitochondrial function and promote cell survival upon stress and aging. One of the MSR pathways is the mitochondrial unfolded protein response (UPRmt), which is well-characterized in *C. elegans* (Quiros et al. 2016; Shpilka and Haynes 2018). The UPRmt is a proteotoxic stress response that senses protein-folding perturbations, which overload the capacities of the mitochondrial quality control network (Jovaisaite et al. 2014). The prototypical UPRmt is best characterized in *C. elegans*; when unfolded or misfolded proteins accumulate in the mitochondrial matrix in the worm, CLPP-1, a mitochondrial protease, cleaves these proteins into small peptides, which are then exported into the cytosol by the mitochondrial inner membrane peptide transporter HAF-1. On the other hand, those released peptides can impede the import of proteins into the mitochondria (Yoneda et al. 2004; Haynes et al. 2010; Naresh and Haynes 2019). The transcription factor ATFS-1 therefore cannot be imported into the mitochondrial matrix where it is degraded by the LON protease (Nargund et al. 2012). As a result, ATFS-1 translocases and accumulates in the nucleus, forming a complex with the small ubiquitin-like protein UBL-5 and the transcription factor DVE-1. Together, this transcription complex activates the expression of nuclear-encoded protein quality components including the heat-shock proteins *hsp-6* and *hsp-60* to re-establish mitochondrial homeostasis (Benedetti et al. 2006; Nargund et al. 2012; Jovaisaite et al. 2014).

Paradoxically, induction of mild mitochondrial perturbations that activate the UPRmt in a moderate fashion have been shown to lead to the extension of lifespan in worms (Dillin et al. 2002; Durieux et al. 2011; Houtkooper et al. 2013), flies (Copeland et al. 2009), and mice (Houtkooper et al. 2013), suggesting that approaches to enhance UPRmt activation may potentially be useful to manage certain age-related diseases. Tetracyclines, such as doxycycline (Dox), are antibiotics that inhibit both bacterial and mitochondrial translation (Houtkooper et al. 2013). Dox has therefore been applied as a pharmacological inducer of the UPRmt (Moullan et al. 2015). Despite evidence that the UPRmt has a beneficial effect on health and aging, the molecular mechanisms that couple the Dox-mediated UPRmt with health- and lifespan extension remain to be elucidated. In addition, these effects of Dox have only been demonstrated in the reference Bristol N2 strain, and the impact of the genetic background on the protective effect of Dox has never been characterized. In this study, we have pharmacologically induced the UPRmt by Dox administration in two genetically divergent worm strains, the N2 and the Hawaii CB4856 strains (Fig. 1A). We show that the Dox-related protective effects are likely mediated at multiple levels of biological regulations, including transcription, translation, lipid, and metabolite levels. Activation of mitochondrial stress response by Dox prolongs lifespan of both worm strains, coupling with some shared and strain-specific features at different layers of regulation. Our findings highlight a strong impact of Dox on mitochondrial functions in both worm strains. Beyond that, Dox administration also upregulated stress response and lipid metabolism, while lowering triglycerides and a comprehensive panel of metabolites in both strains.

**Figure 1.**
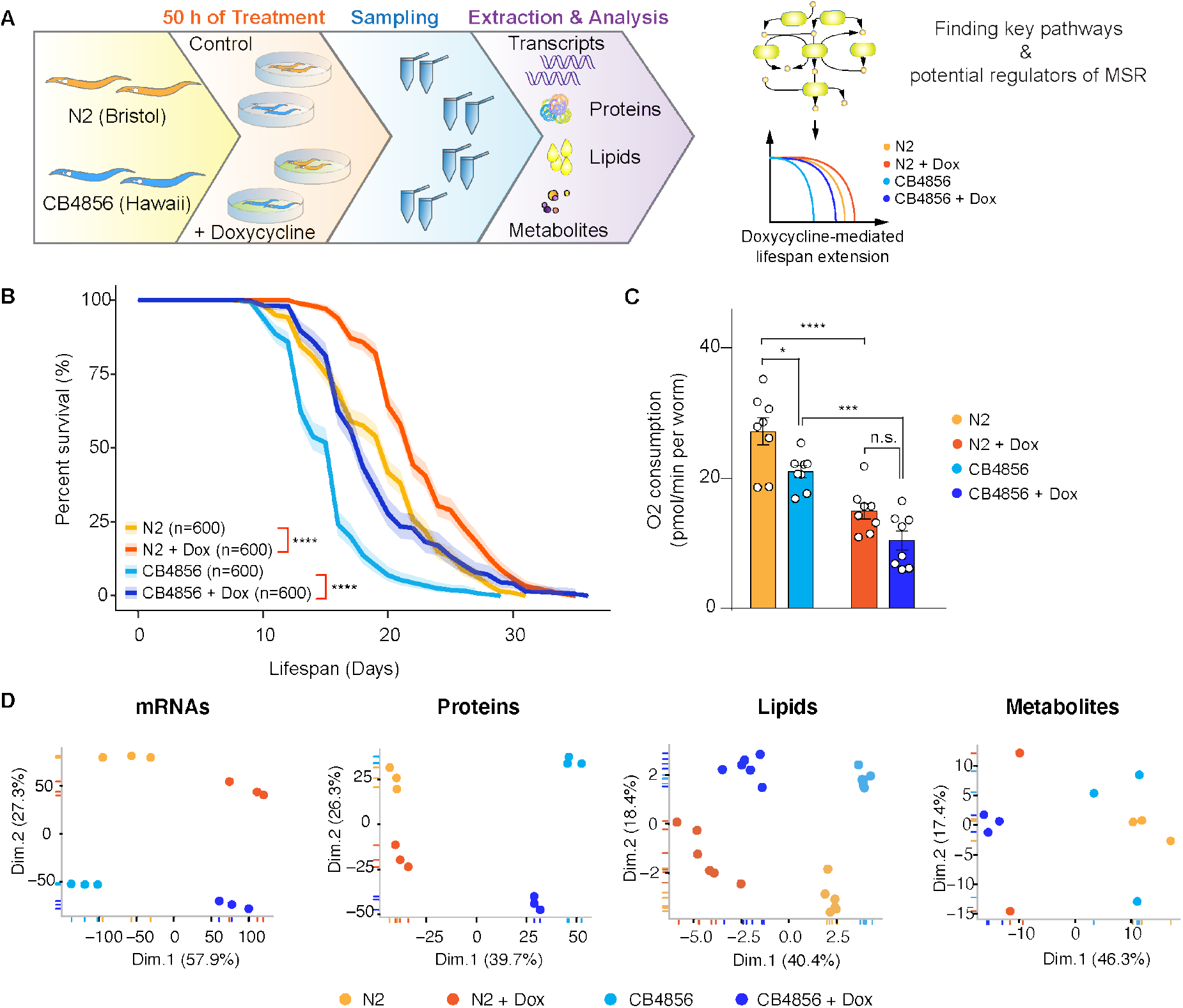
Doxycycline (Dox) activates the mitochondrial stress response (MSR) and prolongs lifespan in two wild-type *C. elegans* strains, N2 (Bristol) and CB4856 (Hawaii). (**A**) Flowchart of the strategy to identify shared and strain-specific MSR regulators. (**B**) Survival analysis of worms fed with control bacteria (*E. coli* HT115) culture on plates with or without Dox (15 μg/mL) by merging nine independent lifespan experiments (n=2400). The shadow area represents the 95% confidence intervals. *P-values* represent comparison with the controls calculated using log-rank test (**: *p*<0.01; ***: *p*<0.001; ****: *p*<0.0001). (**C**) Dox reduced basal and maximal oxygen consumption rate (OCR) in both worm strains. Error bars denote SEM. Statistical analysis was performed by ANOVA followed by Tukey post-hoc test (\**p*<0.05; \*\**p*<0.01; \*\*\**p*<0.001; \*\*\*\**p*<0.0001; N.S., not significant). (**D**) Principal component analysis (PCA) of transcripts/mRNAs, proteins, lipids and metabolites measured in N2 and CB4856 worms treated with or without Dox. **Related to Figure S1, Table S1, S2, S5 and S6.**

## Results

### Doxycycline prolongs lifespan of both N2 and CB4856 strains

To determine whether doxycycline (Dox) has a strain-dependent effect on lifespan, we cultured both N2 and CB4856 worms on plates with or w/o Dox (15 μg/mL) and measured their lifespan in nine independent experiments (Fig. 1B). In basal conditions, CB4856 worms had a shorter lifespan compared to N2 worms. Upon Dox exposure, both worm strains showed an increased lifespan compared to the controls. Because Dox was known to block mitochondrial translation and attenuate respiration (Houtkooper et al. 2013), we measured the oxygen consumption rate (OCR) in N2 and CB4856 worms fed with Dox (Fig. 1C). Respiration was decreased upon Dox addition in both strains. These data suggest that Dox has a universal effect on lifespan and mitochondrial function in both worm strains.

To further investigate the difference in the regulation of Dox-induced longevity in N2 and CB4856 strains, we collected worm samples either exposed to Dox or not, and extracted total RNAs, proteins, lipids and metabolites for multi-omic analysis. We first assessed the strain and Dox effects on each of the omic profiles (Fig. 1D). Interestingly, a primary separation by Dox exposure and a second separation by strain was observed at the transcript and lipid layers (Fig. 1D). Whereas at the protein layer, the first component separated the strains and the second component separated the treatments. For metabolites, we observed a primary separation by Dox followed by a very minor separation by strains. These data suggest that Dox showed a stronger effect on the transcriptomic, lipidomic, and metabolomic layers and the genetic background seems to play a larger role in determining the proteomic layer.

Next, we questioned whether these alterations are commonly shared between the two wild-type strains upon Dox (Supplemental Fig. S1A). In line with the PCA analysis (Fig. 1D), up to 80% of transcripts showed similar Dox-induced changes in both worm strains, in which 2,414 and 2,021 genes were significantly up- and down-regulated, respectively. Consistent with the PCA of the protein profiles, we detected fewer overlapping changes between the two strains upon Dox exposure, in which 127 and 205 proteins were significantly up- and down- regulated, respectively. In particular, the majority of the altered proteins in CB4856 worms were not found to be altered in N2 worms upon Dox (Supplemental Fig. S1A and Supplemental Table S2). Additionally, we also detected a group of genes that showed reciprocal regulation between the two strains, including the 24 at transcript and 65 at the protein level. Lipid and metabolite levels altered similarly in both N2 and CB4856 strains during mitochondrial stress. We then asked how many genes were significantly altered at both transcript and the corresponding protein levels (Supplemental Fig. S1B, Supplemental Table S1 and S2). In N2 worms, 271 transcript-protein pairs were similarly altered in the transcriptomics and proteomics profiles, in which 188 and 83 pairs were up- and down-regulated, respectively, and 66 transcript-protein pairs were reciprocally regulated by Dox. In CB4856 worms, 196 and 181 transcript-protein pairs were up- and down-regulated whereas 172 pairs of mRNAs/proteins were reciprocally regulated by Dox. However, in both worm strains, the genes/proteins that did not change at both transcript and protein levels account for a substantial part of the total altered genes. This reveals that mitochondrial stress induced by Dox may have a global impact on gene expression as well as on post-transcriptional regulation. Collectively, these results suggest that mitochondrial stress induced by Dox prolongs lifespan of both worm strains and the underlying changes at the molecular levels exhibit shared and strain-specific features.

### Defense response and oxidative reduction are affected by genetic variants in CB4856 compared with N2 worms

Since N2 and CB4856 at the basal levels have distinct lifespans as well as different OCR (Fig. 1B-C), we catalogued the genetic differences and their consequences and explored gene and protein expression differences in basal conditions (Fig. 2). We compiled high impact variants present in the CB4856 strain (retrieved from the *Caenorhabditis elegans* Natural Diversity Resource: https://elegansvariation.org/data/release/latest), relative to the N2 reference strain. Among these, we filtered from all the detected variants and selected the major types of consequences from homozygous variants that are of high impact, including frame-shift, start codon lost, stop codon lost, and stop codon gained for further investigations. Across the six chromosomes, most of the detected variants were distributed at the ends of each chromosomes (Fig. 2A and Supplemental Table S3). Of these protein-coding genes, 2,755 had a frameshift (Supplemental Table S3), 32 lost a start codon, 52 lost a stop codon, and 305 had a stop-gained in CB4856 relative to N2 (Fig. 2C).

**Figure 2.**
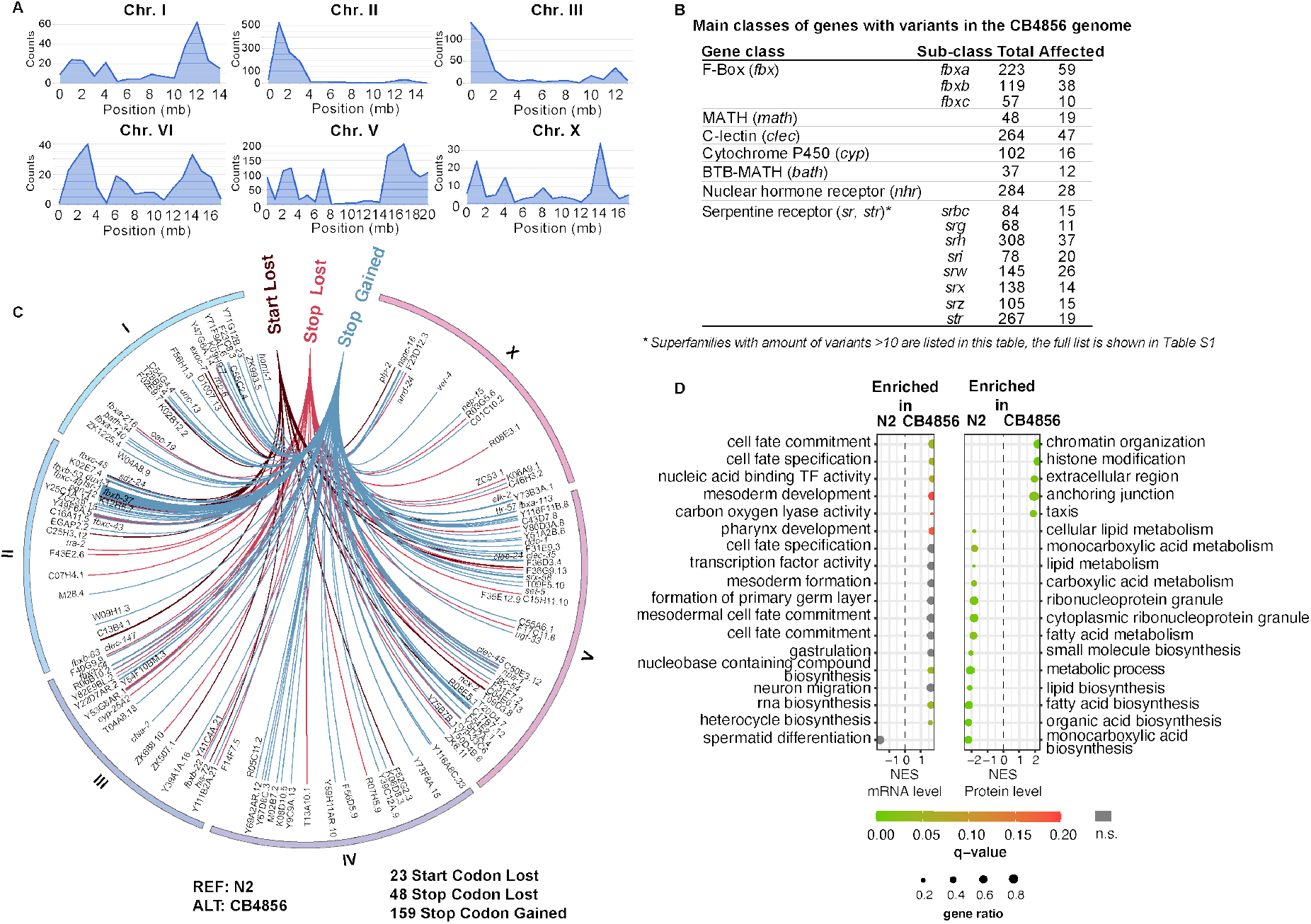
Variant analysis and gene set enrichment analyses (GSEA) determined the differences in the genetic background and gene expression between the N2 and CB4856 strains. (**A**) distribution of high impact variants in CB4856 per chromosome compared to the N2 reference strain. CB4856 VCF (Variant Call Format) file was retrieved from https://elegansvariation.org/data/release/latest. (**B**) Main classes of genes with high impact homozygous variants detected in the CB4856 genome are listed in the table. (**C**) As an example, a circos plot of high impact variants (hard-filtered) present in the CB4856 strain compared the N2 reference strain. Analyzed variants are homozygous and with one of the following consequences: Brown: genes with start codon lost; Dark pink: genes with stop codon lost; Blue: genes with stop codon gained. The full list of variants with a soft filter are shown in Table S1. (**D**) GSEA reveals the differences between the N2 and CB4856 strains at both mRNA and protein expression levels under basal condition. Q-value: false discovery rate adjusted p-values, grey dots: non-significant (n.s.); gene ratio: ratio of found genes within a given gene set. NES: normalized enrichment score. **Related to Figure S2 and Table S3.**

These genes with codon variants could be pseudogenes in CB4856 as a rapid means of adaption (Olson 1999); however, the persistence of these regions may suggest the presence of functional and essential genes. The genes altered by the variants are composed of members of large gene classes, including 107 *fbx* (F-box), 19 *math* (MATH), 47 *clec* (C-lectin), 16 *cyp* (cytochrome P450 family), 12 *bath* (BTB-MATH) genes, 28 *nhr* (nuclear hormone receptor), and 190 *sr/str* (serpentine receptor superfamily) genes (Fig. 2B). F-box family, MATH and BTB-MATH gene families encode ubiquitin-dependent proteosome adapters that were largely involved in targeting foreign proteins for proteolysis during pathogen defense (Thomas 2006). The serpentine receptor superfamily belongs to the G-protein-coupled receptors (GPCRs) that are involved in signal transmission and are playing a vital role in controlling innate immunity against bacterial infections (Nagarathnam et al. 2012; Kaur and Aballay 2020). The variants detected in these genes may lead to defects in pathogen defense in CB4856 worms, which partly could contribute to their shorter lifespan at basal condition compared with the N2 strain. Mammalian C-lectins are carbohydrate-binding proteins that have very narrow ligand specificity, and many C-lectins are involved in innate immune response. Although there are 264 genes encoding C-lectins in *C. elegans*, functions and roles of most C-lectins remain unclear (including the four detected *clec* genes), except for the few of them that were annotated as innate immune genes (Pees et al. 2016). The cytochrome P450 family is composed of enzymes that catalyze oxidative reactions and their substrates include lipids, exogenous and xenobiotic chemicals (Harlow et al. 2018; Herholz et al. 2019). Nuclear hormone receptors are a family of transcription factors that often influence lipid metabolism in *C. elegans* (Chinetti et al. 2000; Ashrafi et al. 2003; Taubert et al. 2006). Variants found in the stop codons of these genes could impair protein function and thus affect lipid oxidation or other oxidation-reduction process. Overall, differences in these gene families could partially explain the lifespan difference between the two wild-type strains under control conditions.

To further explore the impact of the genetic background on the transcript and protein levels, we examined the top gene sets enriched in N2 or CB4856 at the control condition (Fig. 2D and Supplemental Fig. S2). Interestingly, we detected overall more pronounced differences level between the two strains at the protein level, as compared to those detected at the transcript level. At the transcript level, the top enriched gene sets were mainly in CB4856 worms, including those involved in cell fate and transcription factor activity. In contrast, the majority of the top detected gene sets at the protein level were more pronouncedly enriched in N2 worms, including lipid metabolism, carboxylic acid metabolism, ribonucleoprotein granule, metabolic process, and biosynthesis of small molecules, fatty acids, organic acids, and monocarboxylic acids. In CB4856 worms, proteins involved in chromatin organization, histone modification, extracellular region, anchoring junction, and taxis were more enriched, compared to those detected in N2 worms.

Next, we generated enrichment maps for each worm strain to broaden the view and explore other interesting gene sets (Supplemental Fig. S2). Of note, we observed a presence of various gene sets involved in the nucleotide metabolism, and lipid metabolism in the N2 worms at the transcript level, compared to those in CB4856 (Supplemental Fig. S2A). Additionally, a large gene set cluster enriched for lipid metabolism and a cluster of genes enriched for defense response was detected at the protein level in the N2 worms (Supplemental Fig. S2C). Hence, in line with our variant analysis results (Fig. 2), genes involved in defense response, and lipid metabolism were more enriched in N2 worms in control conditions compared with those in CB4856 worms (Supplemental Fig. S2A,C). In the CB4856 worms, we detected a trend of very large gene set for nucleic acid metabolism especially those are associated with RNA metabolism at the mRNA level (Supplemental Fig. S2B). At protein level, we detected a cluster of gene sets enriched for neuron development (Supplemental Fig. S2D), in addition to the gene sets already mentioned above (Fig. 2D). Altogether, these data provide details on strain specificity of the gene set enrichment at both transcriptomic and proteomic level and suggest that some of these differences could be attributed to genetic variants present in the CB4856 genome.

### Dox upregulates lipid metabolism, proteolysis, and stress response in both strains

After observing a considerable impact of genetic variants in the CB4856 strain, we wondered whether the molecular changes upon mitochondrial stress will be affected by the different genetic backgrounds as well. We first analyzed differentially expressed genes using gene ontology (GO) analysis on the 2,414 upregulated transcripts and 127 upregulated proteins in both worm strains upon Dox exposure (Supplemental Fig. S1A, and Fig. 3A,B). At the transcript level, factors involved in proteolysis, lipid metabolism and transport, stress response, innate immune response, and muscle contraction were enriched in both worm strains upon Dox (Fig. 3A). At the protein level, we found fewer GO-terms compared to those at the transcript level, and most of them were related to defense response, including response to stimulus, immune response, response to oxidative stress and to osmotic stress (Fig. 3B). Additionally, we also detected a group of GO-terms related to metabolic processes of flavonoids, organic acids, and monocarboxylic acids.

**Figure 3.**
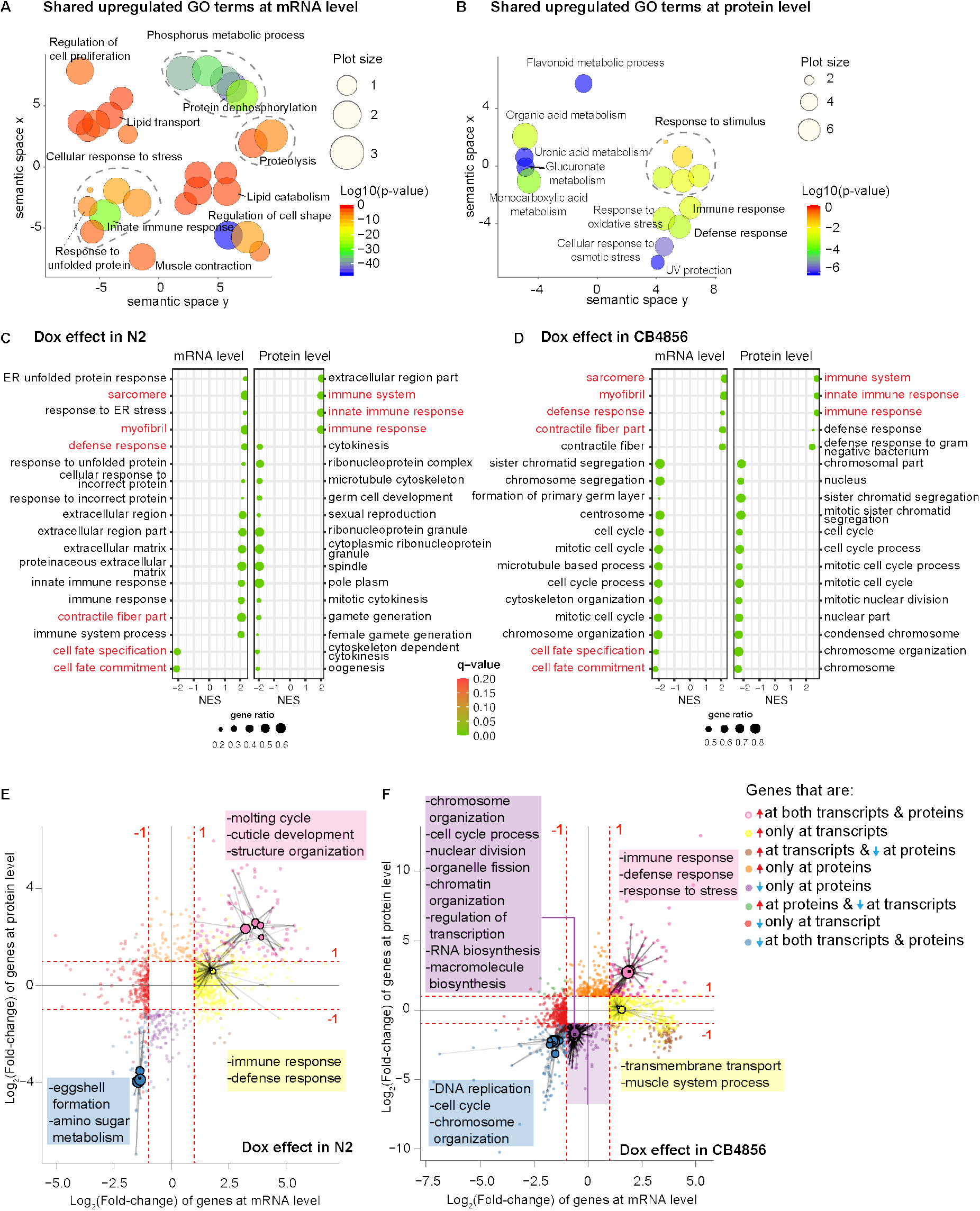
GO-term enrichment and GSEA revealed shared key players and strain-specific regulators of Dox-mediated longevity in N2 and CB4856. (**A**) GO term enrichment analysis (biological process) on the upregulated transcripts shared between N2 and CB4856 upon Dox exposure identified via David and ReviGO. The size of the dots indicates the frequency of the GO term in the underlying Gene Ontology Annotation database; the plots are color-coded according to significance (Log10-transformed); level of significance increases from red to blue. GO terms belonging to the same cluster were grouped and circled in dark grey dashed lines. (**B**) Go term enrichment analysis on upregulated proteins shared by N2 and CB4856 worms upon Dox. (**C-D**) GSEA showed top 18 enriched gene sets in N2 (**C**) and CB4856 (**D**) worms upon Dox at both transcript and protein level. The shared gene sets between the two strains are highlighted in red. Q-value: false discovery rate adjusted *p-values*; gene ratio: ratio of found genes within a given gene set. NES: normalized enrichment score. Shared gene sets are indicated in red. (**E-F**) Scatter plots of differentially expressed genes (*p*<0.05, Log_2_(fold-change)>1 or <−1)) at mRNA (x-axis) and protein level (y-axis) of N2 (**E**) and CB4856 (**F**) upon Dox. Go-term enrichment analysis was then performed on each of the 8 different categories (defined based on directionality change at mRNA & protein levels). Significantly enriched GO terms are presented as a circle (radius determined by gene ratio) and edges to the related genes are shown in grey. **Related to Figure S3**.

Similar to the number of transcripts exclusively altered in either N2 or CB4856 worms upon Dox (Supplemental Fig. S1A), we also detected fewer N2-specific and more CB4856-specific enriched GO terms (Supplemental Fig. S3A,B). In the N2 strain, transcripts related to proteolysis, defense response, carbohydrate metabolism, cuticle development, and oxidation-reduction process were upregulated exclusively upon Dox (Supplemental Fig. S3A). In CB4856, the majority of the GO-term clusters have already been identified in the shared GO-term profile (Fig. 3A), suggesting that CB4856 worms required expression of more genes in order to cope with the mitochondrial stress induced by Dox.

### Dox induces various responses in N2 and CB4856 worms at transcript and protein levels

Next, we determined the top enriched gene sets at mRNA and protein levels separately to assess the effects of Dox on N2 and CB4856 strains. Most top enriched gene sets at the mRNA level in N2 worms were upregulated, while the majority of the top detected gene sets at protein level were downregulated (Fig. 3C). In CB4856 worms, top enriched genes were primarily downregulated at both transcript and protein levels (Fig. 3D). Moreover, immune responses were upregulated in both worm strains upon Dox at the protein level, and gene sets annotated as extracellular region and defense response were upregulated in N2 and CB4856, respectively. Downregulated proteins upon Dox showed strain-specific gene set enrichments, in which proteins related to ribonucleoprotein complex and reproduction were only decreased in N2, whereas those involved in cell cycle and chromosome organization were exclusively decreased in CB4856. Although Dox is known to inhibit mitochondrial translation (Houtkooper et al. 2013), our data confirm that Dox may also attenuate cytosolic translation and this in turn leads to a downregulation of protein sets in different worm strains (D’Amico et al. 2017; Molenaars et al. 2020).

To better understand whether the changes detected at transcriptomic and proteomic levels were representative of one another, we compared the transcriptomic and proteomic profiles of N2 and CB4856 upon Dox (Fig. 3E,F). We divided significantly altered transcripts and proteins (adjusted *p-value*<0.05 & an abs. fold-change>1) into eight categories based on their differences in a co-regulation analysis. An overrepresentation analysis on the detected genes was performed on each of the eight categories to determine the major altered gene sets. In N2 worms, gene sets related to molting cycle, structure organization and cuticle development were co-upregulated at both mRNA and protein level upon Dox (Fig. 3E, pink box). Genes enriched for immune response and defense response were primarily up-regulated at the mRNA level (yellow box). In contrast, genes related to eggshell formation and amino sugar metabolism were co-downregulated at both mRNA and protein levels (blue box). In CB4856 worms, those that encode factors for immune response, defense response, and stress response were co-upregulated upon Dox (Fig. 3F, pink box). Gene sets involved in transmembrane transport and muscle system process were only upregulated at transcriptional level (yellow box). As expected, a number of gene sets were exclusively downregulated at the protein level, including those enriched for chromosome organization, cell cycle, nuclear division, organelle fission, chromatin organization, transcription, biosynthesis of RNA and macromolecules (purple box). Lastly, genes involved in DNA replication, cell cycle and chromosome organization were co-downregulated at both levels (blue box). These data suggest that shared and strain-specific regulation is present at both transcriptomic and proteomic levels upon mitochondrial stress.

### Dox affects mitochondrial gene expressions of both worm strains in a similar fashion

As mitochondria are the target of Dox, we then sought to investigate whether expression of specific groups of mitochondrial genes were altered by Dox in the two strains. About 138 mitochondria-related genes were significantly changed in at least one worm strain upon Dox. We first annotated the altered mRNAs based on their associated functions to assess the effects on the mitochondria (Fig. 4A and Supplemental Table S4). Most genes that encode the mitochondrial complex I, II, and IV, were downregulated upon Dox exposure in both worm strains, whereas those that encode complex III protein remained unchanged. Moreover, we also detected Dox-induced downregulation in transcripts involved in energy-consuming processes, such as DNA replication, DNA repair, apoptosis, mitochondrial translation, and oxidation-reduction process. Many genes involved in transport, such as mitochondrial ATP translocase, *ant-1.3* and *ant-1.4* (Hoshino et al. 2019), and mitochondrial ion transport, *sfxn-1.1*, *sfxn-1.2*, *sfxn-1.3*, *sfxn-1.4*, and *sfxn-5*, were upregulated in both worm strains. For pathways that are involved in energy production, although transcripts involved in the TCA cycles were not changed in the same trend, we detected an overall upregulation in genes involved in mitochondrial fatty acid oxidation (FAO). These data suggest an enhanced energy support from FAO upon Dox.

**Figure 4.**
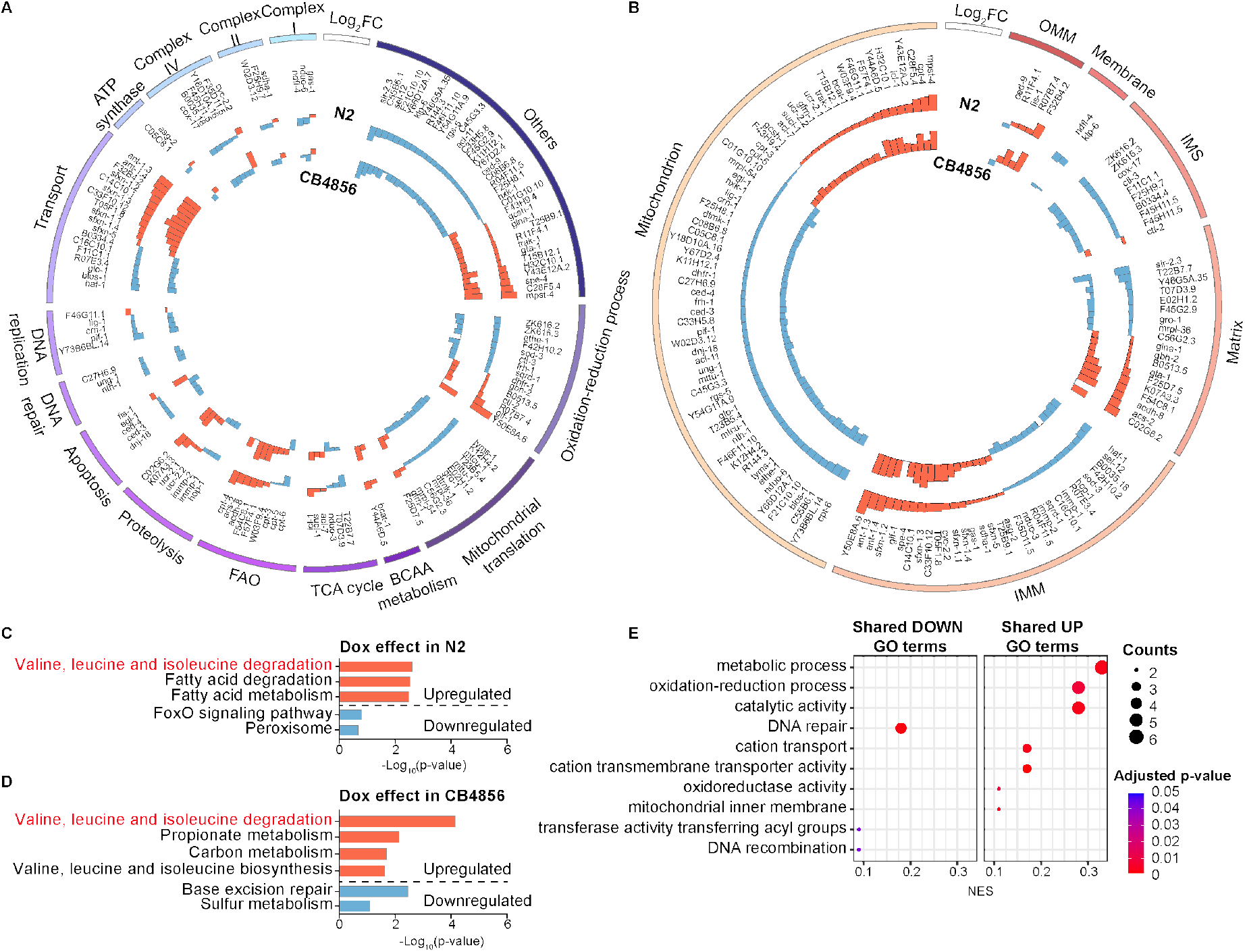
Dox induced significant alterations in mitochondrial genes in both N2 and CB4856 worms. (**A**) Mitochondrial genes associated with different functions are significantly altered in both worm strains upon Dox at the transcript level. Outer ring: N2; inner ring: CB4856. Orange bars: upregulated genes by Dox; blue bars: downregulated genes by Dox. (**B**) Changes in the expression of mitochondrial transcripts encoding for proteins localizing at different mitochondrial compartments after Dox. (**C-D**) KEGG pathway enrichment analysis of up- and down-regulated mitochondrial genes upon Dox in N2 (**C**) and CB4856 (**D**). Orange bars: upregulated genes; blue bars: downregulated genes. Pathway in red: shared between the two strains. (**E**) GO enrichment analysis of the Dox-affected mitochondrial genes in both worm strains. **Related to Figure S4 and Table S4**.

When we plotted the altered mitochondrial transcripts based on their localization within the mitochondria, we noticed that genes that encode proteins/enzymes located in the inter membrane space (IMS) and for membrane proteins were down-regulated, whereas those at outer mitochondrial membrane (OMM) were mainly upregulated in both worm strains (Fig. 4B). Genes that encode for proteins functioning in the mitochondrial matrix and inner mitochondrial membrane (IMM) were partly upregulated and downregulated upon Dox exposure. Because Dox mainly inhibits mitochondrial translation (Houtkooper et al. 2013), we then sought to assess the alterations of mitochondrial proteins upon Dox (Supplemental Fig. S4). However, due to technical challenges in proteomics measurements, we were not able to collect the comprehensive changes at the protein level for these mitochondrial genes upon Dox (only detected 59 mitochondrial proteins were significantly altered).

To determine pathways that are significantly affected by Dox, we performed KEGG pathway and GO-term enrichment analysis on these transcripts (Fig. 4C-4E). In N2 worms, degradation of BCAAs and fatty acids were significantly upregulated whereas FOXO signaling pathway and those associated with peroxisome functions were downregulated by Dox (Fig. 4C). In CB4856 worms, metabolic processes involved in BCAAs, propionate, and carbons were upregulated and those involved in base excision repair and sulfur metabolism were downregulated by Dox (Fig. 4D). We then wondered if there were more similarities in the mitochondrial gene sets shared between the two strains besides BCAA degradation. Under Dox, we detected a number of upregulated GO-terms in both worm strains, including those involved in catalytic activity, oxidation-reduction process, carbon transmembrane transporter activity, cation transport and mitochondrial inner membrane (Fig. 4E). The downregulated mitochondrial genes shared between the two worm strains upon Dox, were mainly enriched for DNA repair, DNA recombination, and transferase activity transferring acyl-groups (Fig. 4E). These GO terms were similar to the ones that we detected in co-downregulated gene sets (Fig. 3E,F), suggesting that the majority of downregulated proteins were related to mitochondria, which could be a consequence of inhibited mitochondrial translation by Dox.

### Dox induces triglyceride degradation by lysosomal lipase to sustain energy production by fatty acid oxidation

In the above-mentioned analysis performed on transcript and protein levels, we detected a large number of genes directly or indirectly linked to lipid metabolism, such as lipid transport, lipid catabolism, and mitochondrial fatty acid metabolism. To expand on the characterization of lipid-related features, we profiled lipids for N2 and CB4856 worms exposed to Dox (Fig. 5, Supplemental Fig. S5, Supplemental Table S5). As the primary component of lipidomic PCA was separated by treatment and the second component was separated by strains (Fig. 1D), we expected to detect more shared and less strain-specific lipidomic changes in the two worm strains. In total we detected 1,572 lipids that belong to 38 lipid classes, which could be categorized further into seven main lipid categories consisting of monoacylglycerol lipids, diacylglycerol lipids, fatty acids/esters, glycerophospholipids, sphingolipids, sterol lipids, and triacylglycerol lipids (Supplemental Fig. S5A).

**Figure 5.**
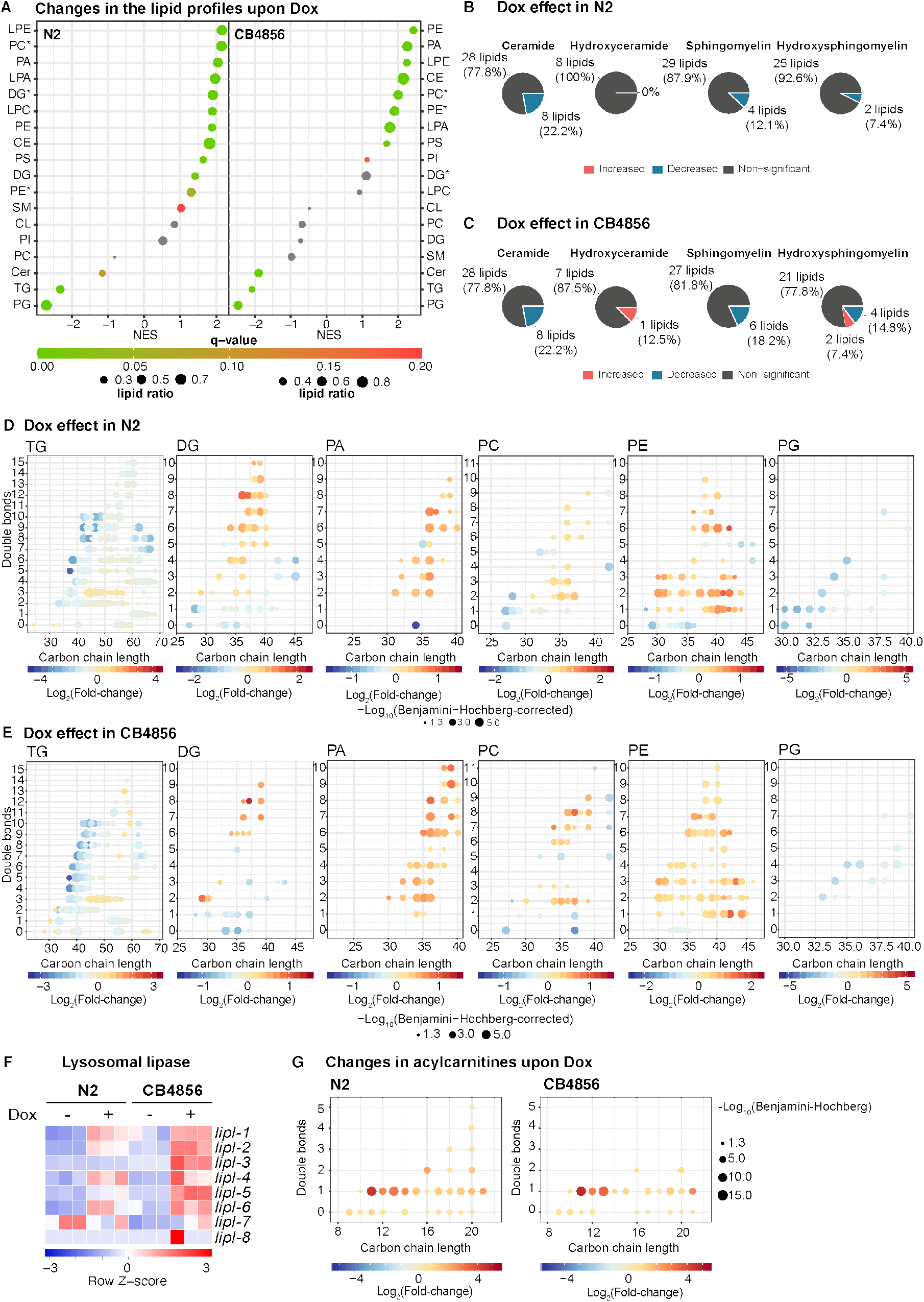
Lipids were significantly altered in Dox-treated worms. (**A**) Lipid class over-representation analysis was performed to identify top changes in the lipidomic layer in N2 and CB4856 worms upon Dox exposure. The asterisk (*): indicates the glycerol position sn-2. (**B-C**) Percentages of increased and decreased lipids in different sphingolipids, including ceramides, hydroxyceramides, sphingomyelins and hydroxy-sphingomyelins in N2 (**B**) and CB4856 (**C**) worms upon Dox exposure (*p<0.05*). A Benjamini-Hochberg corrected *p*<0.05 was applied to determine statistical significance. (**D-E**) Changes in different lipid classes, including triglyceride (TG), diglyceride (DG), phosphatidic acid (PA), phosphatidylcholine (PC), phosphatidylethanolamine (PE) and phosphatidylglycerol (PG) in Dox-treated N2 worms compared to N2 controls (**D**) and in Dox-treated CB4856 compared to CB4856 worms (**E**). (**F**) Lysosomal lipase genes are upregulated at transcript level upon Dox in both N2 and CB4856 worms. (**G**)The majority of acylcarnitines was significantly increased in Dox-treated worms in variant of their genetic background. The dotted-line indicates the significance threshold (*p*<0.01). Y-axis: number of double bonds; x-axis: number of carbon-chain length. **Related to Figure S5 and Table S5.**

We then determined the percentage of lipids showing significant changes in each worm strain upon Dox (Supplemental Fig. S5B,C). Although the overall lipid changes were similar between the two strains, we detected more decreased lipid species exclusively in the N2 worms (126 reduced lipids), compared to those only lowered in CB4856 worms (31 reduced lipids) upon Dox (Supplemental Fig. S1A). In addition, about half of the lowered lipids by Dox are TGs, of which 143 and 105 TGs were decreased in N2 and CB4856 worms, respectively (Supplemental Table S5). To better understand the overall impact on the lipidomic profiles upon Dox, we performed overrepresentation analysis on the lipid classes to identify the top changed lipid species upon Dox in each worm strain (Fig. 5A). Among the top altered lipid classes, most of them increased in N2 worms upon Dox, including LPE, PC, PA, LPA, DG, PLC, PE, CE, PS, DG, SM, CL, and PI. Different than those detected in N2 worms, fewer classes of lipids were increased in CB4856 worms upon Dox, including PE, PA, LPE, CE, PC, LPA, PS, and PI (Fig. 5A). Intriguingly, three classes of lipids, Cer, TG and PG were lowered in both worms upon Dox (Fig. 5A).

Notably, the sphingolipids, such as ceramides, hydroxyceramides, sphingomyelins, and hydroxy-sphingomyelins were significantly altered upon Dox; many of them decreased in both strains except few hydroxy species which were increased exclusively in CB4856 worms (Fig. 5B,C). Low levels of sphingolipids, especially low ceramides, have been shown to couple with increased lifespan in *C. elegans* (Cutler et al. 2014; Huang et al. 2014; Kim et al. 2016) and with caloric restriction in mouse liver (Green et al. 2017). It is therefore not surprising that these sphingolipids were decreased in both worm strains upon Dox exposure, which universally increased lifespan.

In addition, we also observed some shared features in other lipid species, including increased level of DG, PA, PC, and PE, and decreased level of TG and PG (Fig. 5A). Although some TGs with three double bonds slightly increased in both worm strains and alkyl-diacylglycerol (TG[O]) increased in N2, the majority of the significantly altered TGs were decreased upon Dox (Fig. 5D,E). In line with this, we also detected a marked increase of DGs with >2 double bonds and decrease of those with ≤1 double bond, irrespective of the carbon chain lengths in Dox-treated worms. This could be due to either enhanced lipolysis of TGs or reduced TG synthesis. To assess this hypothesis, we checked the changes in the expression of genes encode the enzymes that are involved in both TG synthesis and breakdown pathways. The transcripts of genes involved in TG synthesis, *i.e.* diglycerol acyltransferase *dgat-2* and *mboa-2* remained unchanged in both worms upon Dox (Supplemental Table S1), whereas transcripts of genes involved in TG breakdown, belonging to both adipocyte triglyceride lipase ATGL family (Srinivasan 2015) and lysosomal lipase family, *lipl-1*, was upregulated in both worms upon Dox (Fig. 5F). The transcript of another gene that belongs to the ATGL family, *lipl-3*, was upregulated exclusively in CB4856 worms upon Dox. In addition, transcripts of many other lysosomal lipase genes, including *lipl-2*, *lipl-4*, *lipl-5*, and *lipl-6*, were higher in CB4856 worms, and *lipl-4*, and *lipl-6* in N2 worms upon Dox (Fig. 5F). However, in the proteomics analysis, *lipl-2*, *lipl-5* and *lipl-7* abundances were not significantly changed upon Dox (Supplemental Table S2), suggesting that the regulation of the ATGL family enzymes occurred mainly at the transcriptional level.

We then analyzed the fatty acid and acylcarnitine profiles (Supplemental Fig. S5D,E, and Fig. 5G). In both N2 and CB4856 worms, fatty acids remained largely unchanged upon Dox, except for increases of two polyunsaturated fatty acids (PUFAs), C18:3 and C20:4, and a decrease of saturated fatty acid (SFA) C22:0 in N2, and increases of C18:0, C18:1 and C15:1 in CB4856 worms (Supplemental Fig. S5D). Strikingly, we detected a major change in the acylcarnitines upon Dox in both strains (Fig. 5F), with levels of medium-chain and long-chain acylcarnitines being largely increased in response to mitochondrial stress. In line with the detection of upregulated FAO transcripts in both worms upon Dox (Fig. 4A-D), we therefore speculated that FAO was activated during mitochondrial stress induced by Dox, to produce energy using fatty acids that were released from TGs by lipolysis. Consistent with this, CPT-1 protein was significantly upregulated in both strains with Dox treatment (Supplemental Table S2).

### Dox reduces many metabolites in both N2 and CB4856 strains

As we observed that a large number of metabolic pathways and processes were significantly altered at both transcript and protein levels upon Dox, we also questioned whether metabolites involved in these pathways exhibit shared and strain specific variations. Intriguingly, the majority of metabolites (70 out of 79 significantly altered metabolites in N2, 72 out of 80 significantly altered metabolites in CB4856) altered by Dox were significantly reduced in both N2 and CB4856 (Fig. 6A). Among these metabolites, we detected decreased levels of TCA cycle intermediates, such as citrate and fumarate. Amino acids that could fuel the TCA cycle were also decreased, suggesting an overall reduction in TCA cycle activity.

To better understand the overall pronounced decrease in the metabolite profiles, we assigned these metabolites to the cellular components and organelles (Fig. 6B). Not surprisingly, the most affected organelle in both worm strains was the mitochondria. Other organelles that were likely affected or associated with the reduced metabolites in both strains were the nucleus, ER, peroxisomes, Golgi apparatus, and lysosomes. Based on the annotated chemical classes of the reduced metabolites, pathways involved in nucleotide metabolism, such as pyrimidine metabolism, which is partly carried out in the mitochondria (Evans and Guy 2004; Loffler et al. 2005; Lane and Fan 2015), explain the pronounced impact on nuclear metabolites. In line with the reduced metabolic activity upon Dox, these data suggest that nucleotide metabolism and amino acid metabolism were likely slowed down in both strains, and energy production through TCA cycle, seem also attenuated in order to cope with mitochondrial stress.

**Figure 6.**
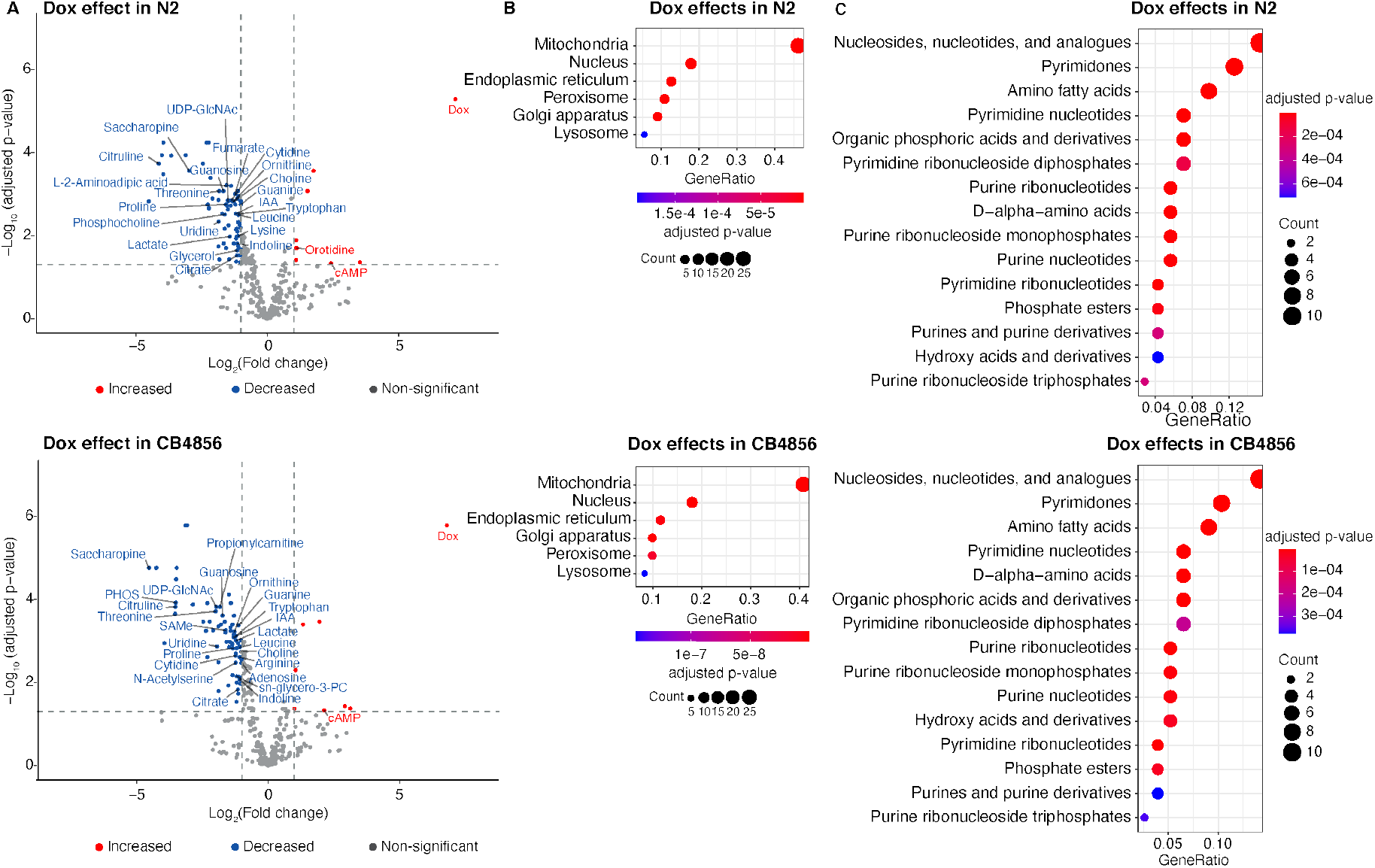
Dox reduces metabolite levels in both N2 and CB4856 worms. (**A**) Volcano plots of the log2 fold change for metabolites (x-axis) against the −Log_10_ adjusted p-value (y-axis) in N2 (top panel) and CB4856 (lower panel) worms upon Dox. (**B**) Cellular localization enrichment analysis of the downregulated metabolites (annotated using the Human Metabolome Database IDs (HMDBs)) in N2 and CB4856 worms. (**C**) Chemical classes enrichment analysis of the downregulated metabolites (annotated using the HMDBs) in N2 and CB4856 worms. For panel **B**and **C**, the size of the dots indicates the number of metabolites for that group; the plots are color-coded according to significance (adjusted p-value); level of significance increases from red to blue. **Related to Table S6.**

## Discussion

In this study, we employed a comprehensive systems approach to explore effects of Dox-induced mitochondrial stress on multiple layers of biology and elucidate the molecular changes in two wild-type *C*. *elegans* strains, *i.e.* N2 (Bristol) and CB4856 (Hawaii). This included the integrated analyses of transcripts, proteins, lipids and metabolites collected in these two worm strains either not exposed or exposed to Dox. Through this approach, we uncovered shared and strain-specific effects of Dox, and important pathways/regulators that are involved in Dox-induced mitochondrial stress response and lifespan extension. Previously, we have showed the robust activation of UPRmt by administering Dox to N2 worms (Houtkooper et al. 2013; Li et al. 2021). It remains, however, unknown whether Dox could also activate the UPRmt and prolong lifespan in other *C. elegans* strains. Although CB4856 worms are genetically divergent and have a shorter lifespan compared with the N2 strain under normal conditions (Doroszuk et al. 2009; Dingley et al. 2014; Thompson et al. 2015; Banse et al. 2019), we observed a comparable effect of Dox on the lifespan and respiratory rate in both worm strains. These data suggest a general and beneficial effect of Dox in terms of lifespan extension. Given the inherent complexity of certain phenotypes, such as lifespan, we assessed whether Dox exhibited shared or strain specific effect at different molecular levels. Based on the PCA of each molecular layer, we noticed that both strain and treatment effects could explain the majority of the difference at each layer of biological regulation.

To better understand the influence of the genetic background in response to mitochondrial stress, we first explored the genetic variants between the two strains in the control condition. In total, we catalogued 3,658 variants with potential disruptive impact on the protein translation, that are distributed on six chromosomes of CB4856 compared to the N2 reference. This number is comparable to the scope of variations detected between these two strains in prior studies (Thompson et al. 2015; Kim et al. 2019). Genes with these homozygous variants mostly belonged to the F-box, MATH, BTB-MATH, C-lectins, Cytochrome P450, serpentine receptor, and nuclear hormone receptor families. Variants that were detected in these families of genes may lead to impaired defense response, oxidative reduction, and lipid metabolism in CB4856 worms. This may partly contribute to their shorter lifespan compared to N2 worms since all these pathways are closely involved in mediation of lifespan (Goudeau et al. 2011; Kaur and Aballay 2020). At the protein levels, we detected more enriched gene sets in N2 worms for lipid metabolism and defense response, supporting our findings in the variant analysis. In addition, we detected more significantly changed mRNAs/proteins in our study compared with a previous study (Kamkina et al. 2016).

Difference in the genetic backgrounds of the two worm strains not only influenced molecular changes at the unstressed condition but also played a vital role upon mitochondrial stress. When analyzing either transcriptomic or proteomic data, we observed a number of shared and strain-specific regulations of different pathways and gene sets. Genes involved in lipid metabolism, proteolysis and stress response were upregulated in both worms upon Dox at the transcript level, which is consistent with the results of previous studies activating UPRmt via other inducers in N2 worms (Liu et al. 2014; Pellegrino et al. 2014; Sorrentino et al. 2017; Liu et al. 2020; Li et al. 2021). Of note, after we integrated transcriptome and proteome profiles, we observed more strain-specific alterations upon Dox. In N2 worms, immune and defense response-associated genes were exclusively upregulated at transcriptomic level, while these genes were upregulated at both transcriptomic and proteomic levels in CB4856 worms. Upregulation of these genes at both transcriptional and translational levels might be a part of a compensatory mechanism balancing the defects in gene expression caused by the large number of disruptive variants present in the CB4856 compared to the N2 worms. In addition, downregulated proteins also showed to be strain-specific, in which ribonucleoprotein complex and reproduction-related proteins were exclusively decreased in N2, and cell cycle and chromosome organization associated proteins were exclusively decreased in CB4856. Our results validated that blocking mitochondrial translation reduces cytosolic translation and leads to a detection of reduced level of proteins in both N2 and CB4856 worms as a consequence (D’Amico et al. 2017; Molenaars et al. 2020).

As the primary molecular target of Dox is the mitochondrial ribosome and mitochondrial translation (Houtkooper et al. 2013), we took a deeper look at the differentially expression mitochondrial genes at both mRNA level and protein level according to their function and localization within the mitochondria. At transcript level, we observed a common effect of Dox on genes involved in various mitochondrial functions. Overall energy-consuming pathways were attenuated, including mitochondrial translation (Houtkooper et al. 2013), consistent with reported observations in worms with mitochondrial stress induced by other stressors (Nargund et al. 2015; Melber and Haynes 2018; Li et al. 2021). Moreover, we observed a metabolic remodeling upon Dox in both worm strains, in which FAO genes were significantly upregulated in both worm strains, suggesting a switch to lipid breakdown to fuel energy production. This speculation was further supported by the upregulated transcripts that belong to the ATGL and lysosomal lipase families, and by the lowered TGs as well as the increased acylcarnitines levels in both worm strains upon Dox. Evidence for enhanced FAO in mitochondrially stressed worms has been documented by others (Baruah et al. 2014; Liu et al. 2020; Tharyan et al. 2020). Moreover, increased expression of fatty acid β-oxidation gene *acs-2* (*fatty acyl-CoA synthetase*, (Van Gilst et al. 2005)) in mitochondrial stressed worms was previously shown to be *atfs-1*-dependent (Wu et al. 2018), suggesting a specific requirement of FAO upon mitochondrial stress. Although upregulated lipid metabolism at the transcript level was not mirrored at the proteomic level, the lipidomic layers still underwent significant alterations in worms exposed to Dox. Besides TGs, sphingolipids, and PGs were significantly decreased in both strains. This was accompanied with an increase of DG, PA, PC, PE.

In our study, we revealed an overall decrease of metabolites in both worm strains upon mitochondrial stress, suggesting a global attenuated metabolism by Dox. The majority of these lowered metabolites were found to be annotated as mitochondrial metabolites, including amino acids, nucleotides and their derivatives, substrates required for TCA cycles. This result correlates well with the lowered expression of mitochondrial genes detected in these pathways. Notably, the lowered nucleotides upon Dox occur concomitantly with the downregulation of transcripts that are involved in DNA replication and repair in both strains. The progression of the replication machinery requires the polymerization of nucleosides triphosphate. As such, low abundance of nucleotides can limit DNA synthesis and arrest the replication process. Likewise, decrease in purine and pyrimidine intermediates may attenuate DNA replication and nucleoside triphosphate synthesis, which could hence prevent the energy consumption in DNA replication.

Collectively, our comprehensive multi-omics systems approach revealed shared and strain-specific pathways that are important for mitochondrial stress-induced longevity in two wild-type worm strains. Effects of Dox-induced mitochondrial stress at different omics layers in the two different worm strains showed more shared features, suggesting universal benefits of Dox. Our data provide new mechanistic insights in Dox-induced mitochondrial stress, which is accompanied by a metabolic rewiring that shuts down anabolic processes and favors catabolic processes including lipid metabolism and TG degradation. This highlights the importance of lipid metabolism in mitochondrial stress-mediated longevity in genetically divergent worm strains.

## Materials and methods

### C. elegans strains

The Bristol strain (N2) and Hawaii strain (CB4856) were used as the wild-type strains obtained from the Caenorhabditis Genetics Center (CGC; Minneapolis, MN). Worms were cultured and maintained at 20 °C and fed with *E. coli* OP50 on Nematode Growth Media (NGM) plates unless otherwise indicated.

### Lifespan measurements

Worm lifespan was performed as described previously (Mouchiroud et al. 2011). In brief, 5-10 L4 worms of each strain were transferred onto RNAi plates (containing 2 mM IPTG and 25 mg/mL carbenicillin) or RNAi plates containing 15 μg/mL Doxycycline (Dox, Cat. D9891, Sigma) seeded with *E. coli* HT115 bacteria. After the progenies reached the last larval stage L4, worms were then transferred onto RNAi (or RNAi + 15 μg/mL Dox) plates containing 10 μM 5FU. Approximately 80 worms were used for each condition and scored every other day. 9 independent lifespan experiments were performed.

### Sample collection for RNA-seq, proteomics, lipidomics and metabolomics analyses

Worms of each strain were cultured on plates seeded with *E. coli* OP50, then eggs were obtained by alkaline hypochlorite treatment of gravid adults. A synchronized L1 population was obtained by incubating the egg suspension in M9 butter overnight at room temperature. Approximately 2000 L1 worms were transferred onto plates with or without 15 μg/mL Dox seeded with *E. coli* HT115. L4 worms were harvested after 50 h by three times of washing with M9 buffer. Tubes containing worm pellets were immediately submerged in liquid nitrogen for snap freezing and stored at −80°C until use. Three biological replicates were collected for each condition.

### RNA extraction and RNA-seq data analysis

On the day of the extraction, 1 mL of TriPure Isolation Reagent was added to each tube. The samples were then frozen and thawed quickly eight times with liquid nitrogen and 37 °C water bath to rupture cell membranes. RNA was then extracted by using a column-based kit from Macherey-Nagel. RNA-seq was performed by BGI with the BGISEQ-500 platform. RNA-seq data analysis was performed using the R package as described previously (Merkwirth et al. 2016). FastQC was used to verify the quality of the mapping (de Sena Brandine and Smith 2019). No low-quality reads were present and no trimming was needed. Alignment was performed against worm genome (WBcel235 ce11 primary assembly and Ensembl release 89 annotation) following the STAR (version 2.73a) manual guidelines (Dobin et al. 2013). The obtained STAR gene-counts for each alignment were analyzed for differentially expressed genes using the R package edgeR (version 3.24.3) using a generalized-linear model (Robinson et al. 2010). The trimmed mean of M values (TMM) method was chosen to normalize the counts and the Cox-Reid common dispersion method for computing the dispersion parameter (tag wise dispersion was also computed). A principal component analysis was also generated to explore the primary variation in the data (Lê et al. 2008; Risso et al. 2014).

### Protein extraction and proteomics analysis

Pellets containing ~2,000 worms were resuspended in 900 μL lysis buffer A (6M Guanidine hydrochloride, 100 mM Tris pH 8.0), and one large stainless-steel bead was added to each 2 mL tube. Tubes were shaken in a bead miller (Retsch) at 30 Hz continuously for 10 min. Then the lysates were placed in a sand bath and heated at 95°C for 10 min. Protein concentrations of lysates were determined using protein BCA assay (Pierce). To extract proteins, 100 μL of lysate were added to 900 μL 100% methanol. Tubes were then spun for 7 min at 15,000 rpm, supernatants were discarded, and protein pellets were allowed to briefly air dry. 50 μL lysis buffer B (8 M urea, 100 mM Tris, 40 mM TCEP, and 10 mM 2-chloroacetamide) were added to the pellets, and the tubes were vigorously vortexed for 10 min. LysC (Wako) was added in 1:50 enzyme: protein ratio, and samples were digested overnight at ambient temperature on a rocker (Fisher Scientific). On the next day, digestion mixtures were diluted down to 1.5 M urea with addition of 50 mM Tris, pH 8.0. Sequencing grade trypsin (Promega) was added in 1:50 enzyme: protein ratio for further 3 h digestion. Samples were then acidified to pH of ~2 by addition of 10% TFA and desalted using Strata SPE columns (Phenomenex). Desalted peptides were lyophilized to dryness in a SpeeVac (Thermo), then resuspended in 50 μL 0.2% formic acid. Final peptide concentrations were determined using Peptide Colorimetric Assay (Pierce). 1 μg peptides were injected onto an in-house high pressure packed capillary column (Shishkova et al. 2018), housed at 55 °C, and separated using Dionex Ultimate 3000 nano HPLC system (Thermo Fisher) over 120 min gradient at 325 nL/min. Mobile phase A consisted of 0.2% formic acid in water, and mobile phase B consisted of 0.2% formic acid in 70% acetonitrile. Eluting peptides were electro-sprayed into and analyzed on Orbitrap Fusion Lumos mass spectrometer (Thermo Fisher). Orbitrap MS1 scans were collected at resolution of 240,000 at 200 m/z with an AGC target of 1×10^6^ ions and maximum injection time set to 50 ms. Precursors were isolated in a quadrupole with the isolation window of 0.7 Th. Tandem MS scans were collected in the ion trap using rapid scan rate, AGC target of 1×10^4^ ions, HCD fragmentation with NCE of 25, and dynamic exclusion of 20 s. RAW files were searched using MaxQuant (Cox et al. 2014) against Uniprot database of *C. elegans*, containing canonical sequences and isoforms. Unless specified, default settings were used. Protein abundances were quantified using label-free quantification with count of 1 and match-between-runs enabled. MS2 tolerance was set to 0.27 Da. Similar differential analysis was performed as for the transcriptome layer using the R package edgeR (version 3.24.3) with a generalized-linear model, similar normalization and principal component analysis methods (Robinson et al. 2010).

### Lipid extraction and data analyses

Around ~2500 worms were collected in a 2 mL tube and the following amounts of internal standards dissolved in 1:1 (v/v) methanol:chloroform were added to each sample: BMP(14:0)_2_ (0.2 nmol), CL(14:0)_4_ (0.1 nmol), LPA(14:0) (0.1 nmol), LPC(14:0) (0.5 nmol), LPE(14:0) (0.1 nmol), LPG(14:0) (0.02 nmol), PA(14:0)_2_ (0.5 nmol), PC(14:0)_2_ (0.2 nmol), PE(14:0)_2_ (0.5 nmol), PG(14:0)_2_ (0.1 nmol), PS(14:0)_2_ (5 nmol), ceramide phosphocholine SM(d18:1/12:0) (2 nmol) (Avanti Polar Lipid). A steel bead and 1:1 (v/v) methanol:chloroform was added to each sample to a volume of 1.5 mL. Samples were homogenized using a TissueLyser II (Qiagen) for 5 min at 30 Hz and centrifuged for 10 min at 20,000 *g*. The supernatant was transferred to a 4 mL glass vial and evaporated under a stream of nitrogen at 45°C. The lipids were dissolved in 150 μL of 1:1 (v/v) chloroform:methanol and lipidomics analysis was performed as described (Molenaars et al. 2021).

In brief, the HPLC system consisted of a Thermo Fisher Scientific Ultimate 3000 binary UPLC coupled to a Q Exactive Plus Orbitrap mass spectrometer using Nitrogen as the nebulizing gas. The column temperature was maintained at 25 °C. For normal-phase separation, 2 μL lipid extract was injected onto a Phenomenex^®^ LUNA silica, 250×2 mm, 5 μm 100 Åand for reverse phase 5 μL of each sample was injected onto a Waters HSS T3 column (150×2.1 mm, 1.8 μm particle size). A Q Exactive Plus Orbitrap (Thermo Scientific) mass spectrometer was used in the negative and positive electrospray ionization mode. In both ionization modes, mass spectra of the lipid species were acquired by continuous scanning over the range m/z 150-2000 with a resolution of 280,000 full width at half maximum (FWHM). For quantification of the data, lipids were normalized to corresponding internal standards according to lipid class, as well as total protein content in samples, determined using a Pierce™ BCA Protein Assay Kit.The dataset was processed using an in-house developed semi-automated metabolomics pipeline written in the R programming language (http://www.r-project.org) (Herzog et al. 2016). Similar differential analysis was performed as for the transcriptome and proteome, using the Bioconductor package limma version 3.42.2 (Ritchie et al. 2015), with a generalized-linear model, similar normalization and principal component analysis methods. Results of the statistical tests were corrected for multiple testing using the Benjamini-Hochberg method.

### Metabolomics analysis

#### Metabolite extraction

Around ~2500 worms were collected in a 2 mL tube. All reagents were placed on ice and samples were maintained at ≤4°C during extraction procedure. A metal bead was added to each sample. Next, 500 μL M1 solvent (methyl tert-butyl ether (MTBE):MEOH=3:1, v/v) was added to each tube and vortexed for 2 min. 325 μL M2 solvent (H2O:MEOH=3:1, v/v) were added to each tube. Samples were vortexed briefly. Then samples were flash-freezed in liquid nitrogen and thawed on ice. This step was repeated three times to facilitate cell breakage. Samples were transferred to a bead-beater and shaken at 1/25 s frequency for 5 min, and this process was repeated 3 times with 5 min interval between each cycle. The samples were centrifuged for 10 min at 12,500 g at 4°C. For downstream metabolomic analysis, 200 μL of the aqueous layer (lower phase) was transferred to glass autosampler vials and dried down by vacuum centrifugation. Aqueous extracts were stored in −80 °C freezer until analysis. Extracts were resuspended in 50 μL 1:1 Acetonitrile:Water and vortexed for 20 s prior to analysis by LC-MS.

#### LC-MS metabolomics

Extracted small molecules were separated on a Sequant ZIC®-pHILIC HPLC column (150 mm × 2.1 mm × 5 μm particle size) at 53 °C using the following gradient: 95% mobile phase B from 0-2 min, decreased to 30% B over next 16 min, held at 30% B for 8 min, then increased to 95% B over next 1 min, then held at 95% B for next 8 min. Flow rate was 130 μL/min. For each analysis, 2 μL/sample was injected by autosampler. Mobile phase A consisted of 10 mM ammonium acetate in 10:90 (v/v) LC-MS grade acetonitrile: H2O with 0.1% ammonium hydroxide. Mobile phase B consisted of 10 mM ammonium acetate in 95:5 (v/v) LC-MS grade acetonitrile: H2O with 0.1% ammonium hydroxide. 2 μL sample was loaded onto column.

The LC system (Vanquish Binary Pump, Thermo Scientific) was coupled to a Q Exactive HF Orbitrap mass spectrometer through a heated electrospray ionization (HESI II) source (Thermo Scientific). Source and capillary temperatures were 350 °C, sheath gas flow rate was 45 units, aux gas flow rate was 15 units, sweep gas flow rate was 1 unit, spray voltage was 3.0 kV for both positive and negative modes, and S-lens RF was 50.0 units. The MS was operated in a polarity switching mode; with alternating positive and negative full scan MS and MS2 (Top 10). Full scan MS were acquired at 60K resolution with 1 × 106 automatic gain control (AGC) target, max ion accumulation time of 100 ms, and a scan range of 70-900 m/z. MS2 scans were acquired at 45K resolution with 1 × 105 AGC target, max ion accumulation time of 100 ms, 1.0 m/z isolation window, stepped normalized collision energy (NCE) at 20, 30, 40, and a 30.0 s dynamic exclusion.

#### Metabolomics data analysis

Data processing was done using Compound Discoverer 3.1 (Thermo Scientific) with untargeted metabolomic study workflow. All peaks within 0-35 min retention time and 100 Da to 5000 Da MS1 precursor mass were aggregated into compound groups using a 15 ppm. mass tolerance and 1.0 min retention time tolerance. Peaks were excluded if peak intensity was less than 1 × 105, peak width was greater than 1.0 min, signal-to-noise ratio was less than 1.5, or intensity was < 3-fold greater than blank. For each feature, compound formula was predicted based on the molecular weight with 15 ppm. mass tolerance. Annotation of the compound formula was assigned by searching database Chemspider with molecular weight or formula. Compound identification was carried out by searching the MS2 spectra against online database mzClound (https://www.mzcloud.org/) as well as a local customized MS2 spectrum library. The customized library is composed of 60 in-house acquired standards, 598 spectra of polar compounds generated from Bamba lab, and whole collection of LCMS MS2 spectra (120K entries) from North American Mass Bank (https://mona.fiehnlab.ucdavis.edu/downloads). Identifications were filtered with a match threshold of 60 for mzCloud hits, and threshold of 30 for customized library hits.

We normalized all the measurements of each sample to correct for bias brought by different amounts of starting material. First, we calculated the sum intensity value of all the metabolite features that were detected in each sample. Original intensity values were divided by the sum intensity of each sample. The resultant intensity values were multiplied by the mean of the sum intensities of all the samples. Normalized intensity values were Log2 transformed before downstream analysis.

### Gene set enrichment analysis (GSEA)

We used the clusterProfiler R package to conduct GSEA analysis on gene ontology biological process terms (Yu et al. 2012). We used a minimum gene set size of 10, a maximum gene set size of 500, and performed 10,000 permutations. For the transcriptome data, we used a gene list that is ordered by log_2_(Fold Changes) from the differential expression analysis. In the proteomic analysis, the gene name for each protein were retrieved and the gene list was then ordered based on fold change values of the protein quantity obtained from the differential analysis of the proteomic level. Pairwise comparisons between layers were performed by overlapping the genes that are up or down with an FDR adjusted p-value of 0.1.

### Transcriptomic/proteomic GO Enrichment analysis

All significantly changing genes / proteins (adjusted p-value < 0.05 and an absolute fold change > 1) were split into 8 groups based on the combination of direction of the fold change, includes genes up & down regulated significantly in a single analysis (i.e. transcriptome OR proteomics). An overrepresentation analysis using GO gene sets was performed on each of the eight groups to determine the main – if any – gene sets changing.

### Variant analysis

Variants for the CB4856 strain were retrieved from the “*Caenorhabditis elegans Natural Diversity Resource*” (https://elegansvariation.org/). Soft filtered variant file was retrieved from http://storage.googleapis.com/elegansvariation.org/releases/20200815/variation/WI.20200815.soft-filter.vcf.gz. Hard filtered variant file was retrieved from http://storage.googleapis.com/elegansvariation.org/releases/20200815/variation/WI.20200815.hard-filter.vcf.gz. These variants were then filtered for the CB4856 strain keeping only 1/1 variants with high impact consequence.

## Acknowledgements

We thank the Caenorhabditis Genetics Center for providing the *C. elegans* strains. We thank all team members of the J. Auwerx laboratory for helpful discussions. This work was supported by grants from the École Polytechnique Fédérale de Lausanne, European Research Council (ERC-AdG-787702) and Swiss National Science Foundation (31003A_179435) and a Global Research Laboratory grant from the National Research Foundation of Korea (2017K1A1A2013124). A.W.G. was supported by the Accelerator prize given by the United Mitochondrial Disease Foundation (PF-19-0232). T.Y.L. was supported by the Human Frontier Science Program (LT000731/2018-L).

## Author contributions

AWG, GEA and JA conceived and designed the project. AWG, AL, TYL, and KH performed the experiments. MM and RHH performed lipidomics analysis, YZ, KO, ES, and JJC performed proteomics and metabolomics analysis, AWG, GEA, AL, TYL, KH, MM, and MBS performed the data analysis. JA supervised the work. AWG, GEA, AL and JA wrote the manuscript with comments from all authors.

## Competing fanatical interests

The authors declare no competing interests.

## Data availability

The RNAseq data has been deposited in the National Center for Biotechnology Information Gene Expression Omnibus database (accession number: GSE159228).

---

**Supplemental Figure S1.**
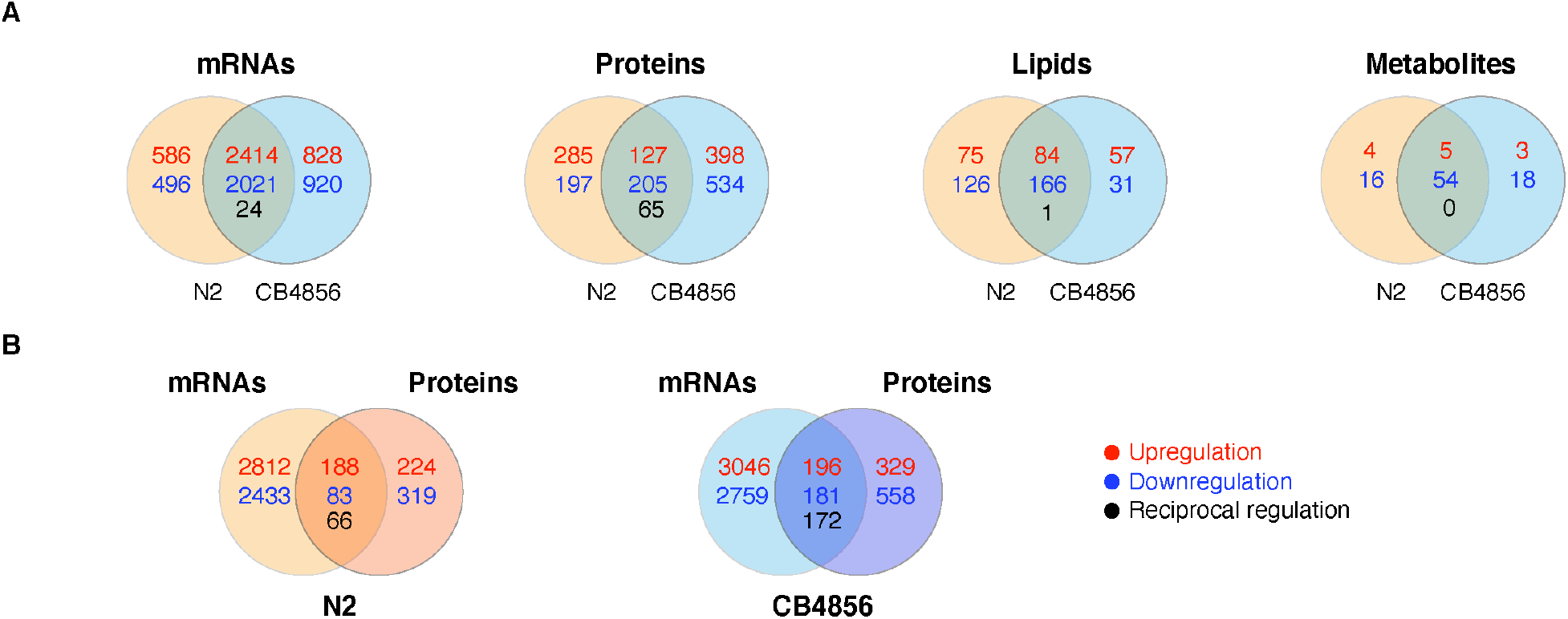
Gene expression changes at mRNA and protein level in both N2 and CB4856 worms upon Dox. Related to Figure 1. (**A**) Venn diagrams depicting the distribution of transcripts/mRNAs, proteins, lipids, and metabolites in N2 and CB4856 worms upon Dox compared to those at control condition. Shared elements (numbers in the middle), N2-specific (numbers on the left side), and CB4856-specific (numbers on the right side) elements in response to Dox treatment are indicated within each pairwise comparison. Upregulation (red), downregulation (blue), and reciprocal regulation (black font) of significantly impacted elements (mRNAs, proteins, lipids and metabolites) are depicted. (**B**) Venn diagrams demonstrate genes significantly altered upon Dox in both N2 and CB4856 worms at mRNA and protein level. Genes that are changed at both mRNA and protein levels (numbers in the middle), only at mRNA level (numbers on the left side), and only at protein level (numbers on the right side) upon Dox treatment are indicated in each pairwise comparison, respectively. Upregulation (red), downregulation (blue), and reciprocal regulation (black) of significantly impacted genes.

**Supplemental Figure S2.**
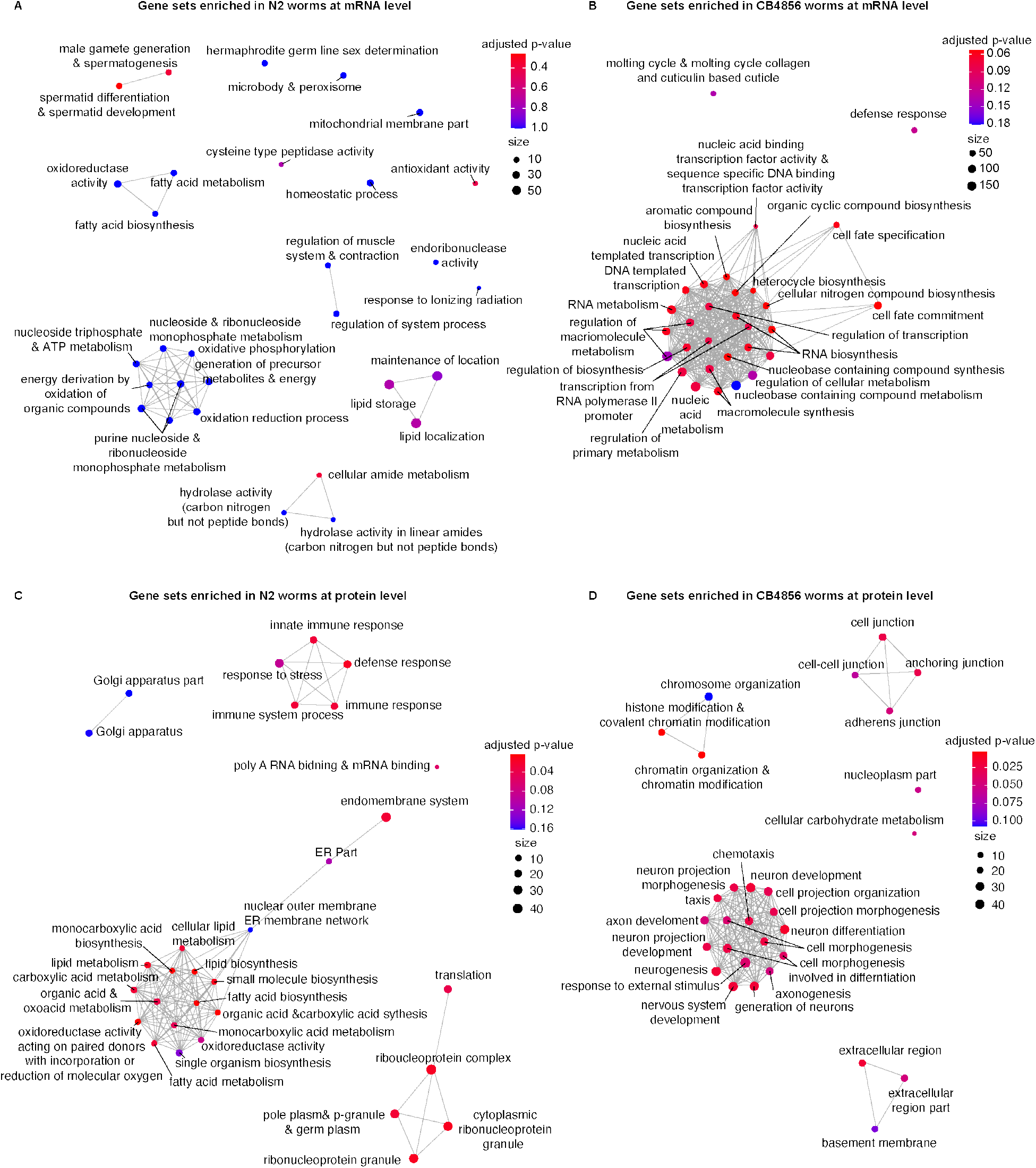
GSEA maps reveal strain-specific differences at transcript and protein level at basal condition. **Related to Figure 2.** The color of the dots represents the adjusted *p-value*; The size of the dots indicates the number of genes within each gene set. The edges represent the presence of mutual genes between the gene sets it connects.

**Supplemental Figure S3.**
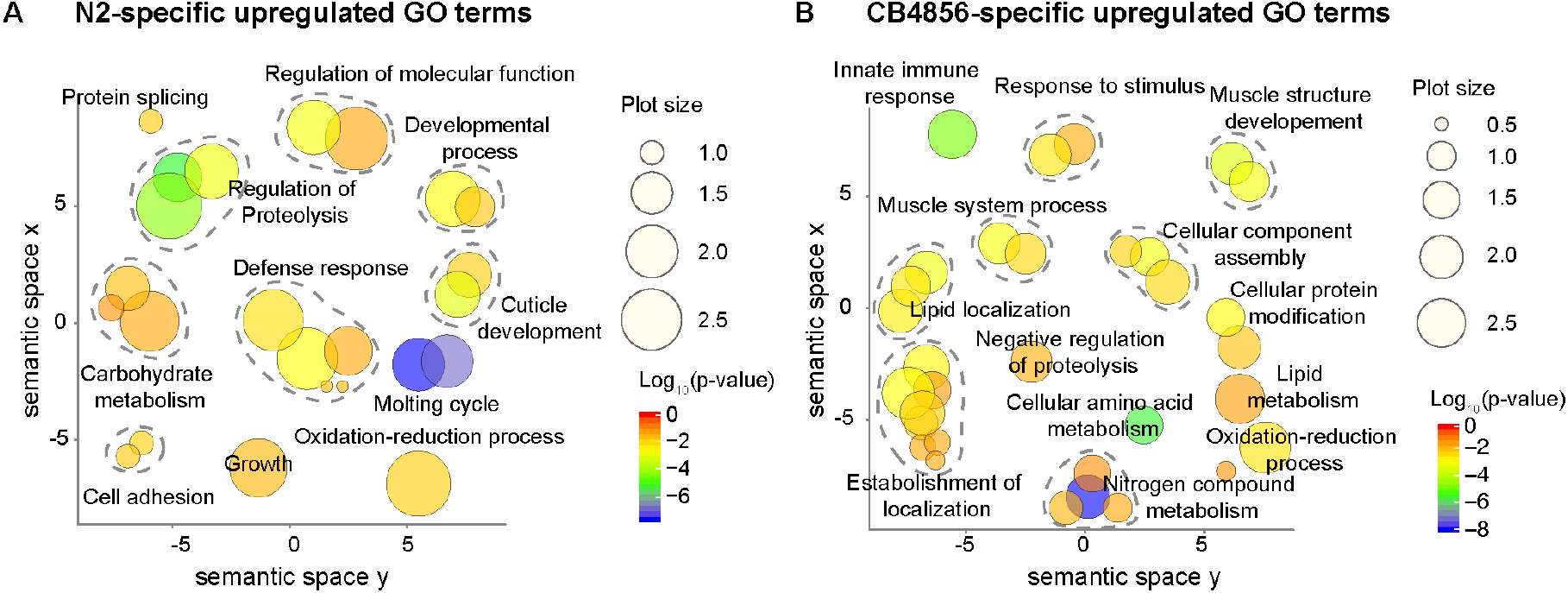
Strain-specific changes at transcriptomic level upon Dox. Related to Figure 3. GO term enrichment analysis was performed on the significantly up-regulated genes exclusively in N2 (**A**), and CB4856 (**B**), respectively. GO term enrichment (biological process) of these genes using David and ReviGO showed upregulated GO terms. The size of the dots indicated the frequency of the GO term in the underlying Gene Ontology Annotation database; the plots are color-coded according to significance (Log10-transformed); level of significance increases from red to blue. GO terms belonging to the same cluster were grouped and circled in dark grey dashed lines.

**Supplemental Figure S4.**
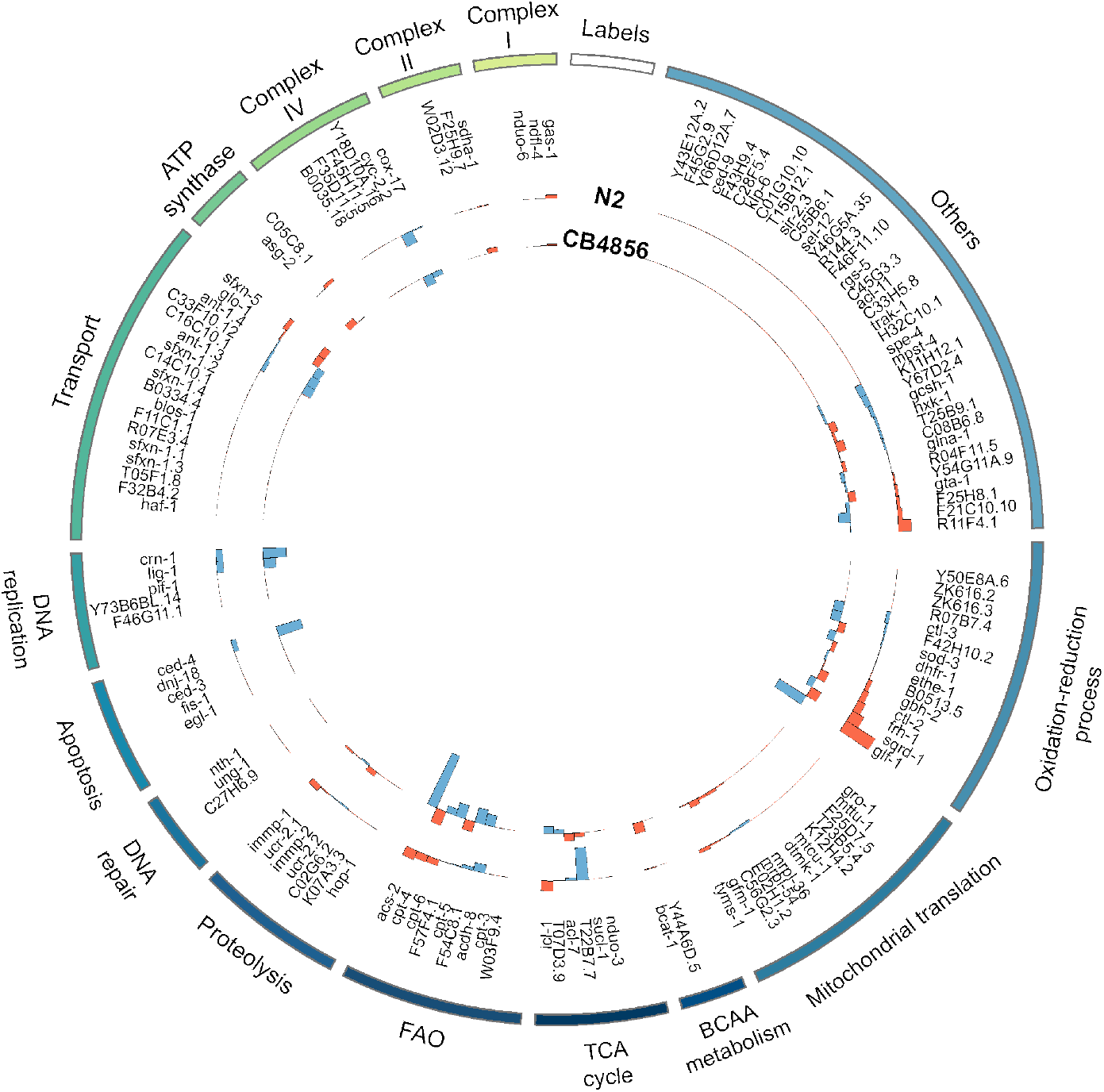
Limits in mitochondrial protein detection revealed less alterations in Dox-treated worms. Related to Figure 4. Changes of mitochondrial proteins are of limited detection. Outer ring: N2; inner ring: CB4856. Orange bars: upregulated genes upon Dox; blue bars: downregulated genes upon Dox. Grey lines: mitochondrial proteins that were not able to detect.

**Supplemental Figure S5.**
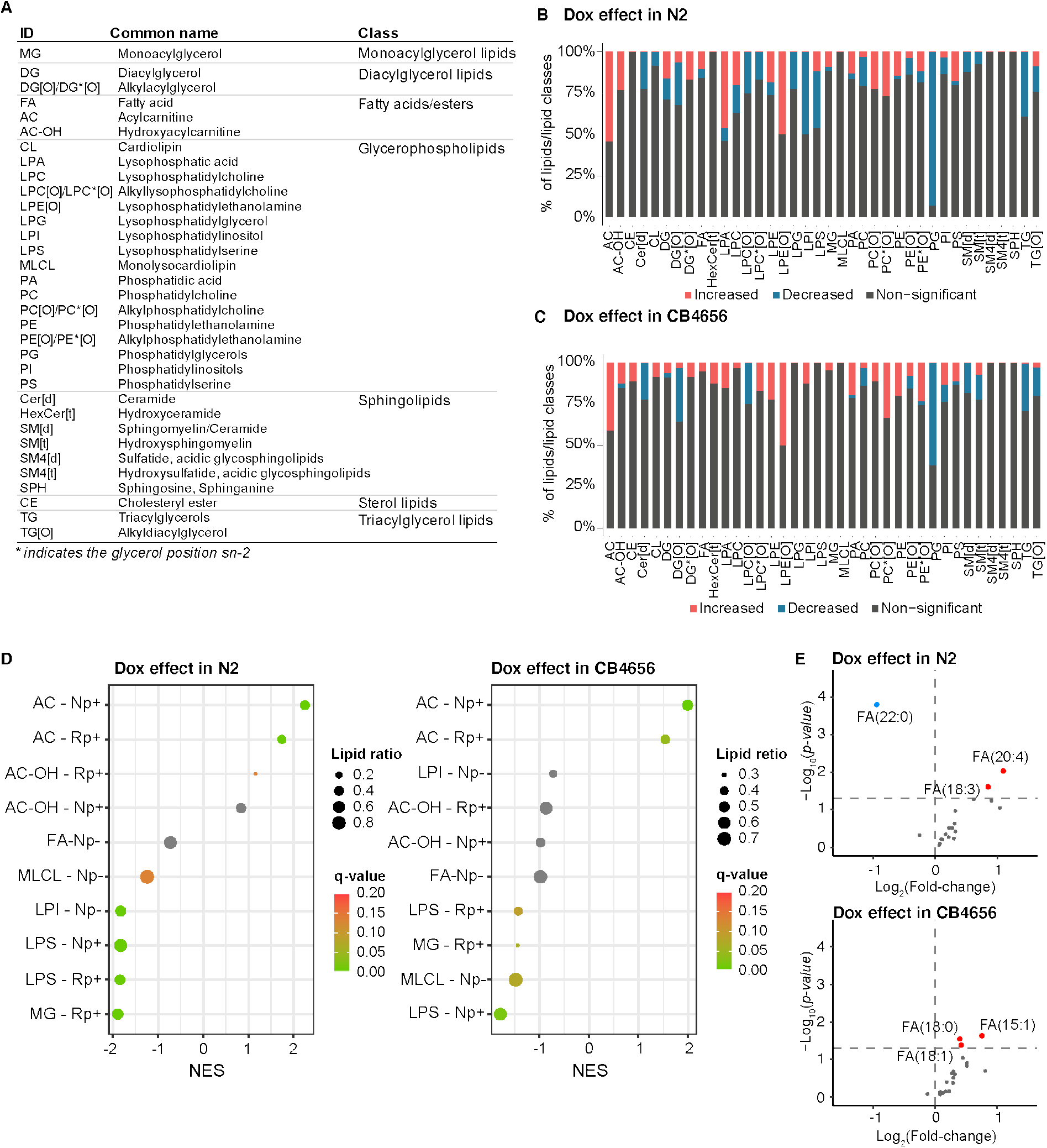
Lipidomics data profiles were significantly altered in Dox-treated worms. Related to Figure 5. (**A**) Overview of measured lipids and their belonging classes. (**B-C**) Percentage distributions of significantly altered lipids species in N2 (**B**) and CB4856 (**C**) worms upon Dox. (**D**) Dox effect in both worm strains on the single chain lipids. Np+: normal phase with positive scan; Np-: normal phase with negative scan; Rp+: reverse phase with positive scan; Rp-: reverse phase with negative scan. (**E**) The majority of fatty acids remained unchanged in Dox-treated worms. The dotted-line indicates the significance threshold (*p-value* <0.05).

